# Cancer patient survival can be accurately parameterized, revealing time-dependent therapeutic effects and doubling the precision of small trials

**DOI:** 10.1101/2021.05.14.442837

**Authors:** Deborah Plana, Geoffrey Fell, Brian M. Alexander, Adam C. Palmer, Peter K. Sorger

**Author notes:** These authors contributed equally. To whom correspondence should be addressed; (cc).

## Abstract

Individual participant data (IPD) from completed oncology clinical trials are a valuable but rarely available source of information. A lack of minable survival distributions has made it difficult to identify factors determining the success and failure of clinical trials and improve trial design. We imputed survival IPD from ∼500 arms of phase III oncology trials (representing ∼220,000 events) and found that they are well fit by a two-parameter Weibull distribution. This makes it possible to use parametric statistics to substantially increase trial precision with small patient cohorts typical of phase I or II trials. For example, a 50-person trial parameterized using Weibull distributions is as precise as a 90-person trial evaluated using traditional statistics. Mining IPD also showed that frequent violations of the proportional hazards assumption, particularly in trials of immune checkpoint inhibitors (ICIs), arise from time-dependent therapeutic effects and hazard ratios. Thus, the duration of ICI trials has an underappreciated impact on the likelihood of their success.

## INTRODUCTION

Extensive effort has been devoted to increasing rates of success in oncology drug discovery by improving preclinical studies (Lindner, 2007; Zhu, 2018; Lin et al., 2019; Hingorani et al., 2019). However, completed clinical trials remain the most valuable single source of information for understanding opportunities and challenges in clinical drug development. Retrospective comparison of specific trials is most commonly performed via meta-analyses and systematic reviews (Haidich, 2010) with the goal of improving patient management in specific disease areas (Whitehead, 2002). Retrospective analysis has also been credited with improving the statistical treatment of trial data, which can be complex and confounded (Weimer and Enck, 2014). However, quantitative analysis of oncology trial results is very difficult to perform at scale because individual participant data (IPD - times of progression, death and censoring), which are necessary for high-quality analysis, are rarely available (Riley et al., 2010; Stewart and Tierney, 2002). Trial results are commonly reported in the form of summary statistics and, in the case of oncology trials, plots of patient survival based on the Kaplan–Meier estimator (Kaplan and Meier, 1958). These plots are generated using IPD but it has proven time-consuming and resource-intensive to access the underlying IPD values because journals and investigators do not generally make them available (Wan et al., 2015).

To address this problem, the International Committee of Medical Journal Editors (ICMJE) recently endorsed the concept of releasing IPD from clinical trials and developed a set of data reporting standards. However, less than 1% of papers published in the last three years actually comply with these standards (Danchev et al., 2021). We and others have therefore developed methods to bypass this problem by using image processing to impute IPD values from published plots of the Kaplan-Meier estimator (Alexander et al., 2018; Guyot et al., 2012; Rahman et al., 2019). Our approach is consistent with the Institute of Medicine’s reports on best practices for sharing data from published clinical trials, including crediting the sources of the data and sharing all code used in the analyses (Committee on Strategies for Responsible Sharing of Clinical Trial Data et al., 2014). In this manuscript we describe a comprehensive analysis of imputed IPD and reconstructed survival curves from ∼150 publications reporting phase III cancer trial results, which in aggregate comprise ∼ 220,000 overall survival or event-free survival events (e.g. progression free survival, PFS).

We find that therapeutic responses as measured by overall survival (OS) or event-free survival fit well to unimodal distributions described by the two-parameter Weibull function; one parameter is proportional to median survival and the second quantifies changes in hazard over time. Analyzing survival functions with parametric distributions has a long history (Boag, 1949), but evidence has been lacking about which distribution to use; our analysis addresses this directly and shows that survival analysis using Weibull forms increases precision without reducing accuracy. For example, we observe that a 50-patient trial (assessing overall survival) is as accurate and precise as a 90-person trial evaluated using traditional nonparametric statistics; this finding is directly applicable to improving power in phase Ib/II trials (Ferrara et al., 2018). Weibull fitting of survival data also confirms that violations of the assumption of proportional hazards are common in contemporary trials (Alexander et al., 2018) notably for immune checkpoint inhibitors but also more broadly. We find that differences in time-varying hazard ratios between treatment arms cause trial duration to impact the likelihood of success. Simulation shows that some failed trials with strong time-dependence might have been judged successful (that is, to confer a hazard ratio <1 at 95% confidence) had they been run for longer. Trial characteristics computed from IPD also make it possible to compare response distributions across diseases and therapeutic modalities, potentially making it possible to improve the design of future trials (Vanderbeek et al., 2019), reduce attrition (Kola and Landis, 2004; Nixon et al., 2017), and assist cost-effectiveness analyses by validating the assumptions necessary for such work in representative patient data (Hoyle and Henley, 2011).

## RESULTS

### Cancer patient survival data can be accurately parameterized across many diseases and drug classes

We used previously described algorithms and approaches (Guyot et al., 2012; Rahman et al., 2019) to mine published papers reporting the results of phase III research clinical trials in breast, colorectal, lung, and prostate cancer with endpoints including OS or surrogates such as PFS, disease-free survival (DFS), and locoregional recurrence (LRR) (which we henceforth consider in aggregate as “event-free survival”). Briefly, image processing was used to extract plots of the Kaplan Meier estimator from figures, while the at-risk tables and the number of patient events were manually extracted from the publication, making it possible to impute IPD (e.g. times of progression, death and censoring; **Fig. 1A**). Trial metadata were also curated and the resulting information is released in its entirety as supplementary materials to this paper and via an online repository, www.cancertrials.io (the accuracy and precision of IPD imputation is discussed in the **Methods;** see also **Supplementary Data Files S1-S2**). Analysis of OS data from 116 figures (from 108 unique randomized controlled trial (RCT) reports) yielded 237 distributions (91,255 patient events) and data on event-free survival from 146 figures (from 135 unique RCT reports) yielded 301 distributions (127,832 patient events). Classes of therapy included chemotherapy, immune checkpoint inhibitors, radiotherapy, surgery, targeted therapy, and placebo/observation.

**Fig 1:**
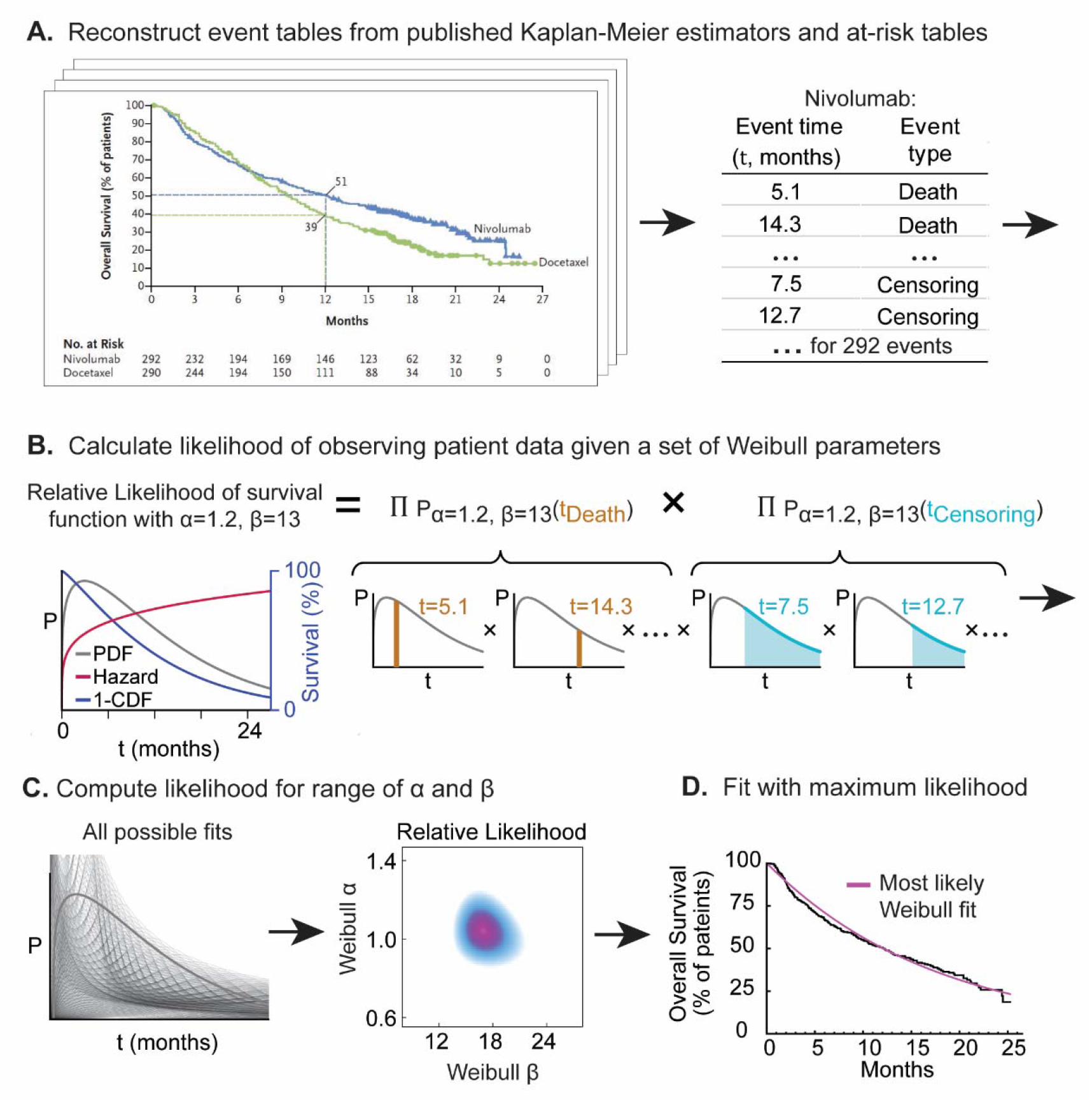
Procedure for parameterizing survival curves starting with published figures. (**A)** Kaplan-Meier survival curve and at-risk table obtained from clinical trial publication. Individual patient data (IPD) were imputed from digitized survival curves and at-risk tables as previously described (see Methods)**. (B)** Each set of parameters corresponds to a different probability density function (PDF) and survival function (which corresponds to 1-the cumulative density function (CDF)). The likelihood of observing actual data is then computed. (**C)** Likelihood calculation is repeated for all possible parameter values. (**D)** The most likely (best) fit is obtained by finding the parameter values with the maximum likelihood.

Multiple parametric forms have been proposed to describe survival in oncology trial data, including the Log Normal, Log Logistic, Gamma, Weibull, Gompertz–Makeham, and Exponential distributions (Caldwell, 2007; Collett, 2003). These differ in their hazard functions, which describe the likelihood of an event (e.g. death or progression) at a given time. Specifically, the exponential distribution assumes a constant hazard, the Weibull, Gamma, and Gompertz–Makeham distributions allow hazards to change monotonically with time, and the Log Normal and Log Logistic functions allow for non-monotone hazard rates (Cox and Oakes, 1988; Kalbfleisch and Prentice, 2002; Klein and Moeschberger, 2003; Lawless, 2003). These distributions differ the most at long follow-up times when their “tails” fall to an asymptotic value or to zero. However, such long event times are rarely recorded in traditional oncology trials, which are limited in duration by cost and increased censoring (often because patients switch to an alternative therapy (Gao et al., 2008)). As a consequence, we found that two types of two-parameter distributions fit survival data equally well: Weibull distributions and Log-Normal distributions (Weibull median R^2^ = 0.981 and Log Normal median R^2^ = 0.980). We selected the two-parameter Weibull distribution for further analysis because its parameters are easily interpreted in terms familiar to oncologists. The Weibull α (shape) parameter describes increasing or decreasing hazard over time (Kleinbaum and Klein, 2012), and the β (scaling) parameter is proportional to median survival time (Matsushita et al., 1992). Survival data fit by Weibull distributions with α <1 have decreasing hazard rates over time, meaning that the likelihood of progression or death is highest at the start of the trial and decreases over time. A value of α =1 corresponds to a constant hazard and α >1 to a hazard that increases with time.

For each trial arm in our data set, we calculated the relative likelihood of different values of α and β to describe its IPD (**Fig. 1B-C**) and found the best-fit parameter values (**Fig. 1D**). It can be helpful to visualize the resulting distributions in three different ways: (i) as a probability density function (PDF), the probability that an event will occur at any point in time *t*; (ii) as a cumulative density function (CDF), the integral of the PDF with respect to *t*; for OS data, 1-CDF is overall survival at *t*; and (iii) as a hazard function, which corresponds to the ratio of the PDF and survival function. A plot of patient-level data as a PDF makes clear that death or progression is right-skewed for all values of α observed here, so that a substantial proportion of all events occur well after the modal (peak) values (as illustrated in **Fig. 1B**). Fitting Weibull distributions therefore quantifies the frequently observed phenomenon that some patients’ response to therapy is substantially better than the most commonly observed responses to that treatment. In oncology trials, the survival function is usually determined using the nonparametric Kaplan-Meier estimator (**Fig. 2A-C**), which accounts for progression or death events as well as censoring (e.g. loss of follow-up within the trial duration or withdrawal from the trial for reasons other than progression or death). Hazard ratios (which measure the treatment effect relative to a control) and their confidence intervals are universally computed using Cox proportional hazards regression (referred to as Cox regression hereafter), a semi-parametric method (Cox, 1972).

**Fig. 2:**
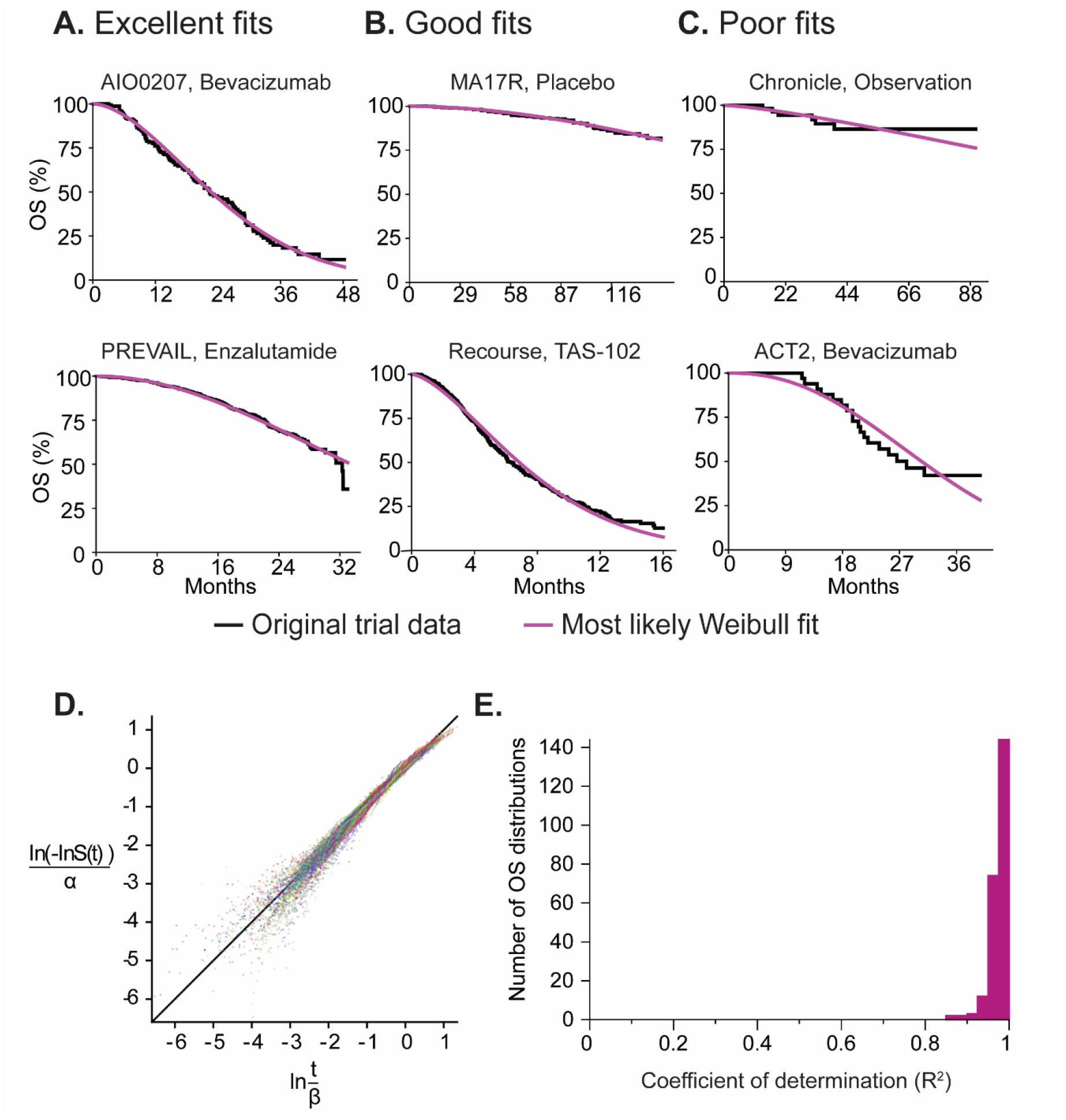
Representative fits of Weibull distributions to overall survival trial data. **(A)** Weibull fits to data for plots of the Kaplan-Meier estimator falling in the top 25^th^ percentile quality of all fits (NCT00973609, NCT01212991) **(B)** at the 50^th^ percentile (NCT00003140/NCT00754845, NCT01607957) and **(C)** in the bottom 25^th^ percentile (NCT00427713, NCT01229813). **(D)** Weibull plot for patient events in all OS trials (for 237 trial arms from 116 publication figures). This is a transformation of survival data that is linear if the data follow a Weibull distribution. **(E)** Goodness of fit as the coefficient of determination (R^2^) explained by fitted Weibull functions for all trial arms reporting OS data.

For trials reporting OS data, we found that a two-parameter Weibull distribution had a median coefficient of determination of R^2^ = 0.981 across 237 trial arms from 116 figures in clinical trial reports (**Fig. 2D, 2E; Methods**). The theoretical maximum R^2^ value can be calculated under the hypothesis that all OS distributions are Weibull distributions and that deviations are attributable only to sample size variability, which gives R^2^ = 0.995. Thus, the error between observation and the Weibull model is ≈1%. For trials reporting event-free survival data (e.g. PFS; **Supplementary Fig. S1**) the median R^2^ was 0.950 as compared to a theoretical maximal R^2^ = 0.996, which corresponds to 5% of observed variance not explained by the Weibull model.

Across the entire data set, the worst fits were observed for trials with relatively few events, for example, the Chronicle trial (NCT00427713 (Glynne-Jones et al., 2014)) with only 7 deaths and 16 progression events in the observation arm **(Fig. 2C; Supplementary Fig. S1**). Fit was also poor for trials involving pre-planned changes in treatment such as the ACT2 trial (NCT01229813) (Hagman et al., 2016), in which treatment induction was followed by randomization to maintenance treatment at 18 weeks (**Fig. 2C**). In these cases, responses varied over the course of the trial by design and a good fit to a single survival function was not expected. For two of the lowest quality fits, a three-parameter Weibull distribution (consisting of the traditional two-parameter distribution with an additional “cure rate” term (Yu et al., 2004)) resulted in an improvement in the quality of fit (**Supplementary Fig. S2**). However, two-parameter forms had excellent performance for the bulk of the data, are generally more computationally tractable for large-scale analysis, and are more parsimonious. The use of a cure rate parameter might be advisable for different sets of data in which cure is a known outcome (e.g. R-CHOP for non-Hodgkin’s lymphoma)(Schmittlutz and Marks, 2021). Biomarker-stratified arms were also described well by a single two-parameter Weibull distribution, as illustrated by the OS and PFS curve fits for panitumumab in combination with FOLFIRI and FOLFIRI alone in wild-type and mutant *KRAS* metastatic colorectal cancer (trial 20050181 (Peeters et al., 2014); **Supplementary Fig. S3A-B**). We conclude that a two-parameter Weibull distribution provides an excellent fit to available trial data across multiple types of cancer, treatment modalities, and metastatic status.

### Deviation from the one-distribution Weibull model in PFS data

The slightly poorer fit of event-free survival data to Weibull distributions as compared to OS data was caused primarily by a steep fall seen in some survival curves at early time points; this behavior has previously been interpreted as evidence for subpopulations of responding and non-responding patients, particularly in trials of immune checkpoint inhibitors (ICI) (Stewart et al., 2017). It has also been attributed to delayed T-cell activation by ICI therapy (Gibson et al., 2017). For trials of these agents, we found that fit to PFS data could be improved by using a mixture model comprising two different two-parameter Weibull distributions each with its own α and β parameters. This is potentially consistent with the two-population hypothesis (**Fig. 3A; Supplementary Fig. S4**). However, the same was also true of control arms in these trials, suggesting that the deviation from a single distribution at early time-points was not ICI-specific. Moreover, a mixture model exhibited no improvement in fit as compared to a single-distribution fit for OS data from ICI or control arms (**Fig. 3B; Supplementary Fig. S4**). When we examined PFS data from an additional 25 ICI trials, we found that the drop in patient survival at early event times (as determined by finding the time *t* corresponding to the greatest change in the slope of the survival curve) was strongly correlated with the time of the first radiological scan (as reported in trials’ methods sections; Pearson correlation 0.982, *p* < 10^-21^; **Supplementary Data File S3**). We then simulated the influence of scan times on PFS by beginning with a single Weibull distribution, and supposing that progression could only be observed when a tumor is scanned, which occurs periodically (often at 9 or 12-week intervals depending on the trial protocol). Accounting for scan time recapitulated the initial decline in PFS in both control and experimental arms of ICI trials, and improved fit to PFS distributions, raising mean R^2^ to 0.98, compared to 0.92 without considering scan times (**Fig. 3C; Supplementary Fig. S4; Methods**). We conclude that the initial drop in PFS arises when observations corresponding to the left-hand tail of a unimodal response distribution are concentrated in time because radiological scans used to measure tumor progression are performed at discrete intervals. Thus, mixture models involving two Weibull curves do not appear necessary to accurately describe survival for ICIs or any other class of therapy that we have examined. Instead, when scan times are accounted for, single two-parameter Weibull distributions are observed to have an outstanding fit (R^2^ =0.98) to PFS data.

**Fig. 3:**
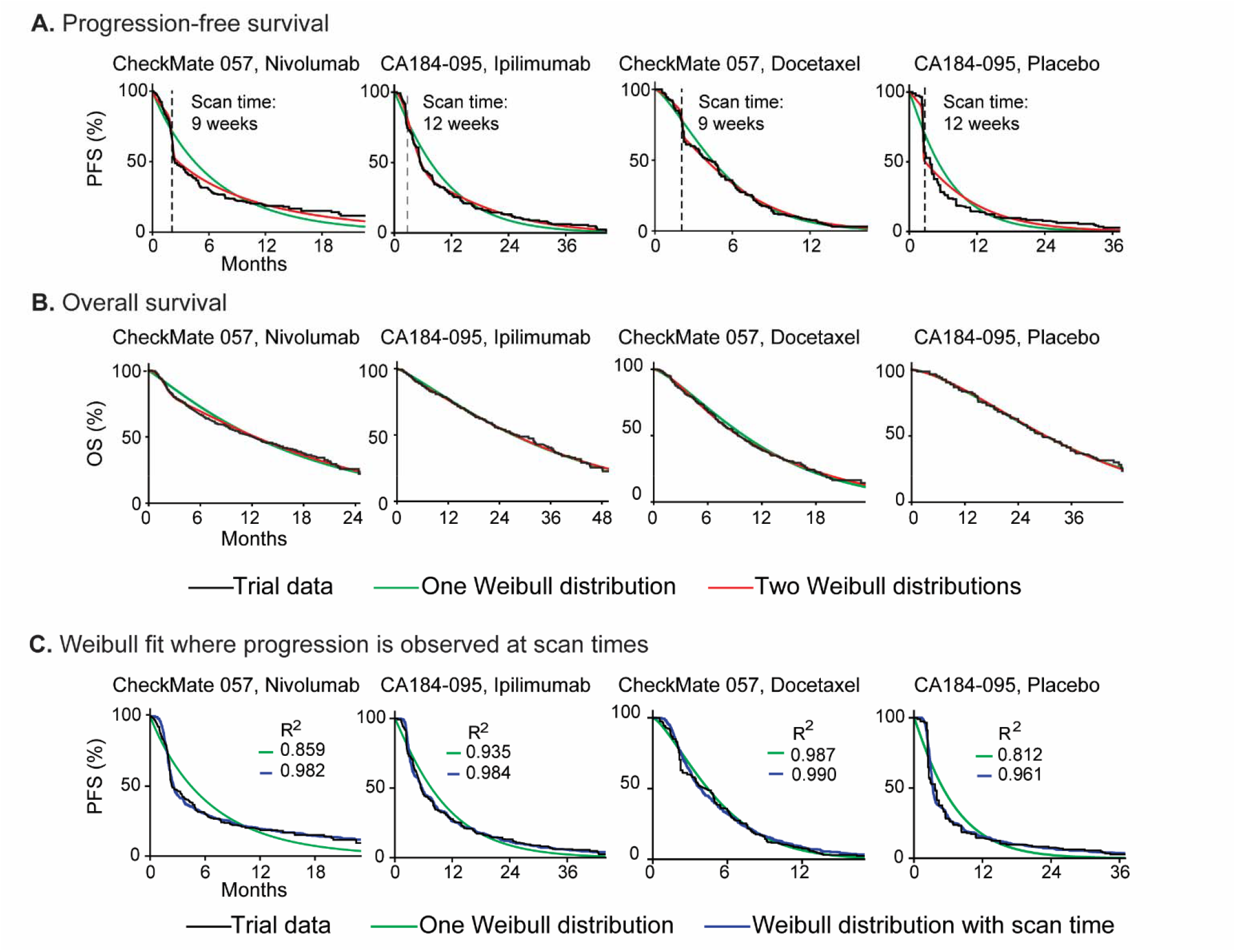
Fit of Weibull models to overall survival and progression-free survival data for trials of immune checkpoint inhibitors. **(A)** Progression-free survival distributions and **(B)** overall survival distributions for individual arms of two trials of immune checkpoint inhibitors (NCT01673867, NCT01057810) with fit to either one or two Weibull distributions. PFS data are best described by using a mixture model of two Weibull distributions, each with two parameters. OS distributions are described equivalently well by a one or two-distribution fit. **(C)** PFS simulations that account for the periodicity of radiological scans to detect progression improve the quality of one-distribution Weibull fits, as quantified by the coefficient of determination (R^2^).

### Parametric fitting improves the precision of drug efficacy estimates

To compare the performance of Weibull-based methods and nonparametric survival estimates, we calculated 12-month pointwise confidence intervals for “small cohorts” simulated by sampling 20 to 100 patients at random from data imputed from real trial arms. Nonparametric methods are unable to report a numeric confidence interval when too few events occur before or after the time point at which the analysis is being conducted (12 months in our simulations; “failed” estimates correspond to 100% or 0% survival respectively). In small cohorts this issue is well known to limit the power of nonparametric analysis. We found that no numerical confidence interval could be computed for 20% to 40% of OS trial arms, and for 23% to 61% of event-free survival trial arms, with higher failure rates occurring with smaller simulation sample sizes. In comparison, use of a Weibull form made it possible to calculate 12-month confidence intervals in every case examined (∼19,000 OS and ∼25,000 event-free survival simulations). The same advantage applies to median survival time, which is relevant because many early-phase oncology trials report median survival (or EFS, PFS) with an estimated upper bound that is “not reached” and therefore uninformative (Cox and Oakes, 1988). Thus, Weibull fitting is broadly applicable in survival analysis of small cohorts because it makes it possible to reliably obtain confidence intervals for median survival and for time points of interest.

For the subset of OS curves in which confidence intervals could be computed using both parametric and nonparametric methods (125 curves), the Weibull-based approach was more precise (it had a narrower confidence interval) across all sample sizes, and accuracy was comparable. By way of illustration, the precision of a 50-patient trial was comparable to that of a 90-patient study using traditional methods (**Fig. 4).** For event-free survival (for which 99 event-free survival curves could be compared), the precision of a Weibull-based approach was also greater than a nonparametric approach across all sample sizes (**Supplementary Fig. S5A**). Thus, modeling cancer patient survival using Weibull functions increases precision, without compromising accuracy. The impact is greatest for trials having sample sizes typical of phase Ib or II trials. This is also a setting in which use of innovative statistical methods is most likely to be acceptable from a regulatory standpoint.

**Fig. 4:**
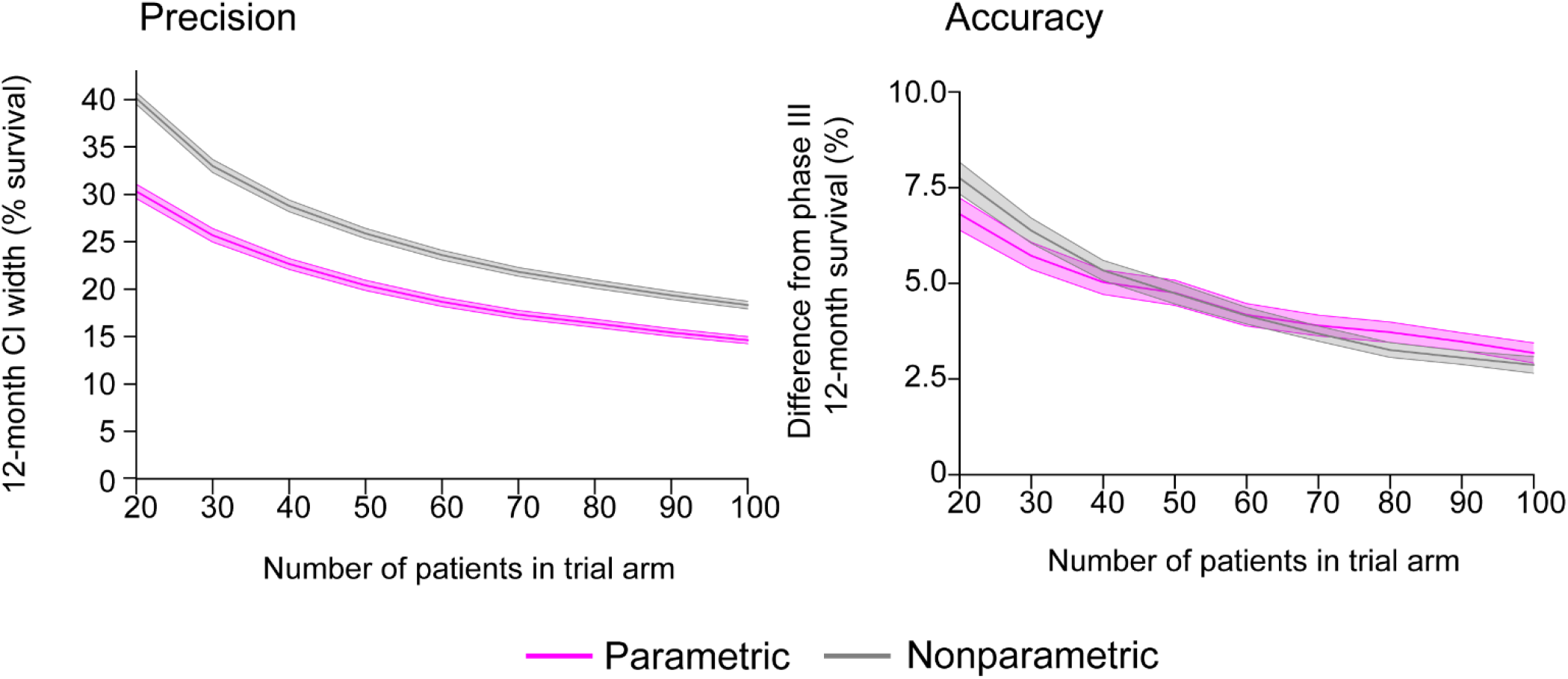
Impact of parametric statistics on precision and accuracy of overall survival confidence interval estimates. Comparison of nonparametric and parametric (Weibull distribution) methods to compute 12-month overall survival confidence intervals for trials with small cohorts (20 to 100 patients) produced by randomly subsampling patient events from 125 actual trial arms. Note that nonparametric estimates did not return a numerical confidence interval for 41% of OS curves, while Weibull fitting made it possible to calculate 12-month confidence intervals for every survival curve in every simulation. Precision is defined as the width of the confidence interval in percent survival. Accuracy is defined as the absolute difference between the 12-month survival estimated from small cohorts (for each of 10 simulations per sample size and trial), and the value computed from all patients in a given phase III trial. Shaded regions are 95% confidence intervals.

As one illustration of the use of Weibull parameterization, we analyzed a recent trial that encompassed both phase I and II data, and tested pembrolizumab with dabrafenib and trametinib for metastatic BRAF-mutant melanoma (MK-3475-022/KEYNOTE-022; NCT02130466) (Ribas et al., 2019). Parametric fitting for 15 patients in phase I yielded a median value of 15.2 months and 95% confidence interval for median PFS of 7.8 to 23 months, while nonparametric estimates yielded a 95% confidence interval of 5.4 months to “not reached” (median value 15.4 months). Nonparametric analysis of a phase II cohort of 60 patients within this same trial demonstrated a median PFS of 16 months and a 95% confidence interval of 8.6–21.5 (Ascierto et al., 2019). Thus, parametric fitting of data from 15 patients made a comparably precise and accurate estimate of median PFS as nonparametric analysis of 60 patients (**Supplementary Fig. S5B**). The availability of more precise inferences would make it possible to use the same number of patients enrolled in single phase II study (e.g.: 60 patients) to perform three different signal-finding studies (each involving 20 patients) with no loss of power. This might have been particularly helpful in the case of KEYNOTE022, a trial which failed to meet its primary endpoint.

### Weibull fitting quantifies differences in survival across cancer types

The availability of a large set of IPD made it possible to search for systematic differences in the parameters of survival distributions by disease class. Best-fit Weibull parameters were compared across cancer types and metastatic status using an ANOVA test with a Bonferroni correction for multiple hypothesis testing at a two-tailed significance level of 0.05 (see **Supplementary Data File S2**). The largest parameter difference was between metastatic and non-metastatic disease, irrespective of tumor type (β values corresponding to median survival of 26 and 174 months respectively). We also observed that β values were significantly larger for breast cancer than lung cancer on a variety of experimental and control treatments (β values corresponding to median survival of 38 versus 15 months in the metastatic setting) **(Fig. 5)**, which is consistent with previous data on relative disease severity (Allemani et al., 2015; National Cancer Institute, SEER.). Event-free survival parameter values were similar to OS values **(Supplementary Fig. S6, Supplementary Data File S2**). Lung cancers had a significantly lower α (shape parameter) for OS as compared to other cancer types (average α =1.30 for lung; versus ∼1.5 to 1.6 for breast, colorectal and prostate cancers), demonstrating a high probability of early death in the course of treatment, even when controlling for differences in median survival times. This difference in shape corresponds to a wider distribution of lung cancer survival times as compared to other cancer types (e.g.: the ESPATUE lung cancer trial had a survival time distribution with a 10^th^ percentile value of 3 months and a 90^th^ percentile value of 49 months; **Fig. 5C**). Parameters drawn from IPD could be used to model cancer survival distributions across diseases, facilitating inter-group comparison in master protocol or basket trials which often involve different cancers types (Palmer et al., 2020; Park et al., 2020).

**Fig. 5:**
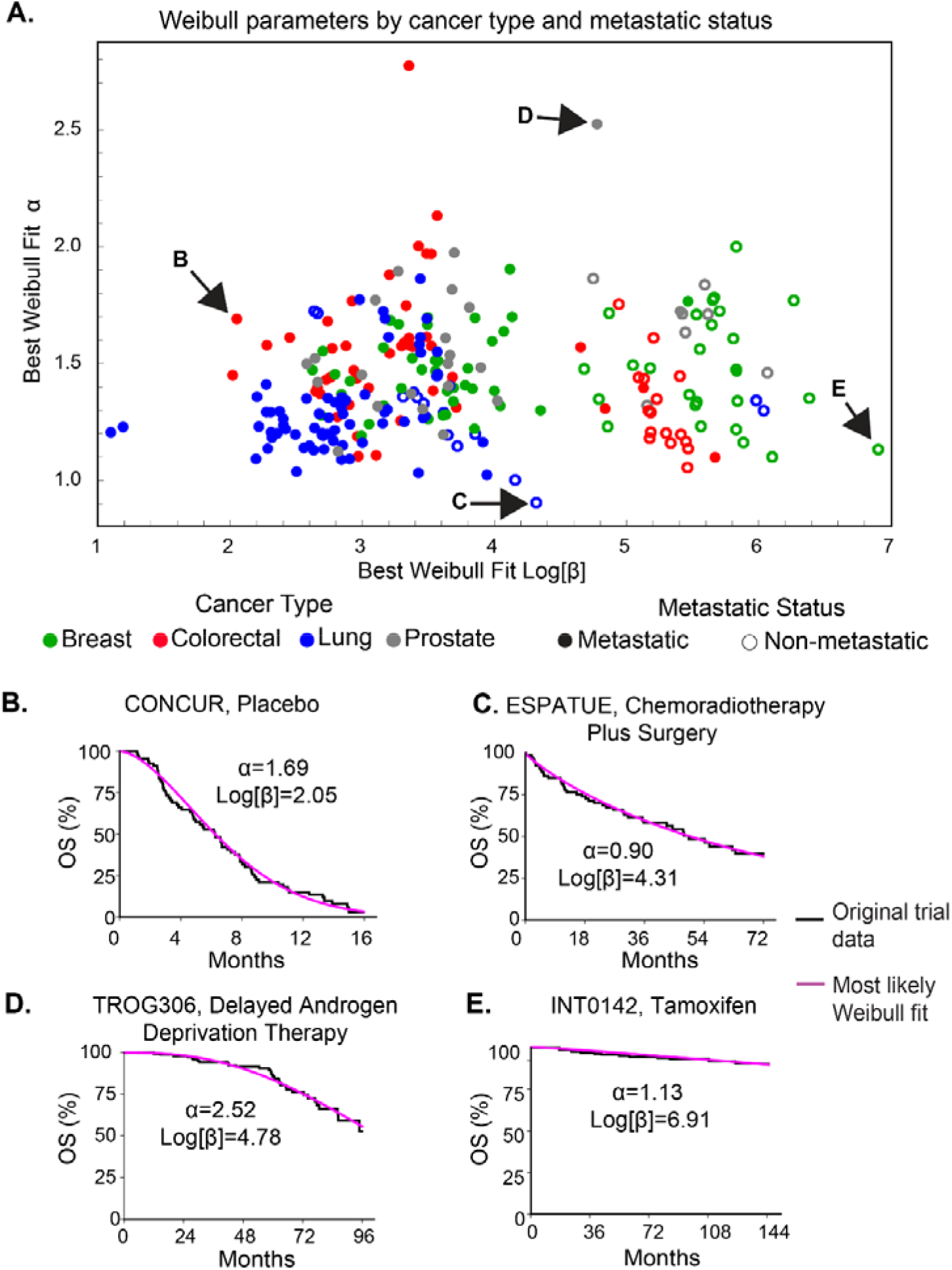
Best fit Weibull parameter values for trials reporting overall survival data. Weibull fits for trials reporting overall survival data. **(A)** Survival distributions categorized by cancer type and metastatic status (defined as trials that included patients with distant metastases). Representative survival functions and fits for trials across a variety of cancer types including **(B)** metastatic colorectal cancer (NCT01584830) **(C)** non-metastatic lung cancer (NCT number not reported**) (D)** metastatic prostate cancer (NCT00110162) and **(E)** non-metastatic breast cancer (NCT number not reported).

### Violations of proportional hazards and impact of trial length on estimates of relative hazard

Randomized controlled trials in oncology are evaluated in a majority of cases based on hazard ratios; if the hazard ratio is significantly below one then the test treatment has decreased the risk of death or progression relative to control, and the trial is regarded as successful (Cox, 1972; Cox and Oakes, 1988). Cox regression estimates the semi-parametric hazard ratio (hereafter referred to as HR_SP_) based on the number and timing of death, progression, or censoring events, all of which increase over time. As expected, when Weibull α and β parameter values were compared between experimental and control arms, a trial was more likely to be successful (HR_SP_ < 1 at 95% confidence) when differences in β values were greater: the median difference between control and experimental β values in OS curves was 0.94% for unsuccessful trials and 29% for successful trials (**Fig. 6A**). A similar pattern was observed for event-free survival data, with control and experimental β values differing by 1.0% for unsuccessful trials and 35.7% for successful trials **(Supplementary Fig. S7**).

**Fig. 6:**
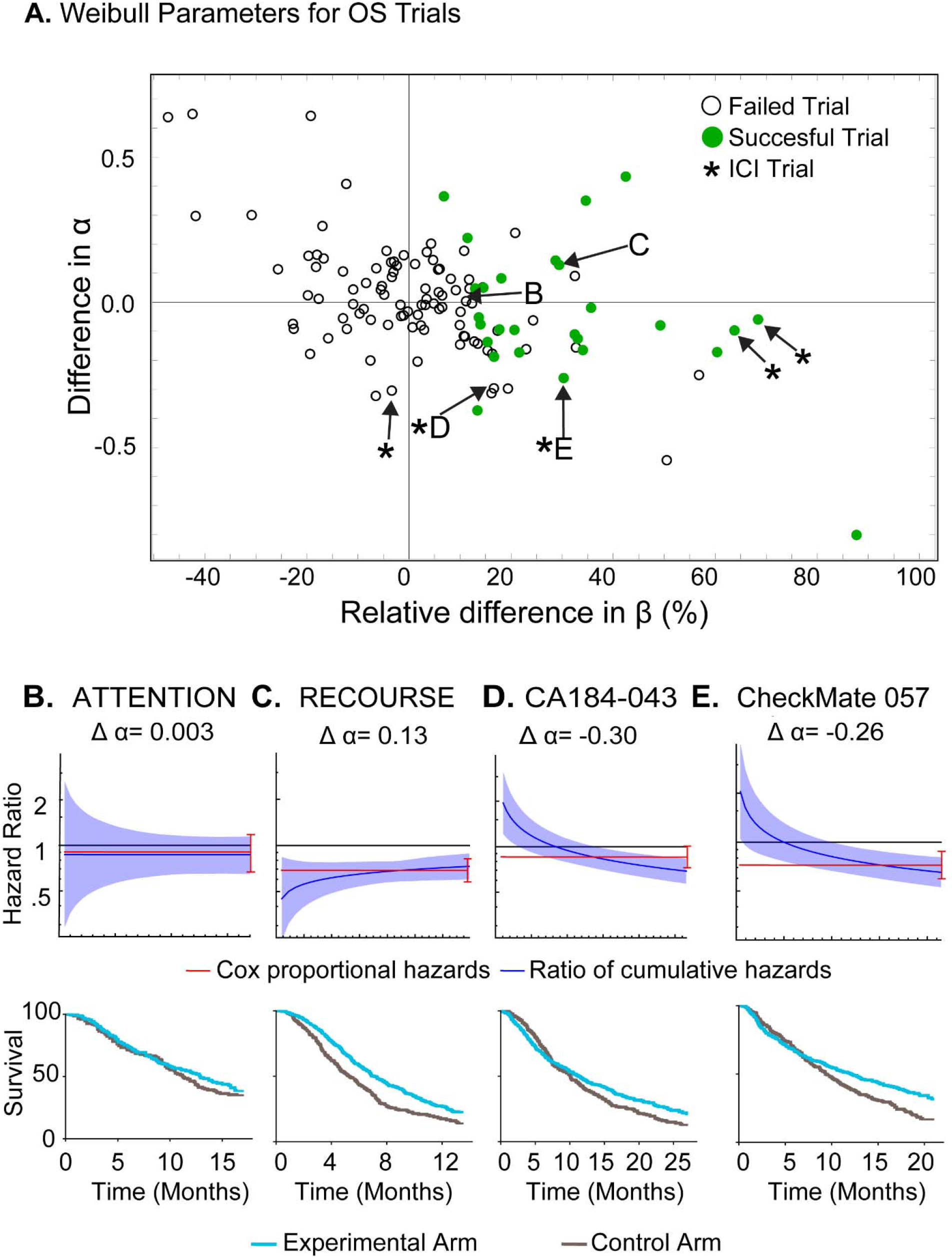
Parameter values for Weibull fits to overall survival data scored by trial outcome. **(A)** Differences in Weibull α and β parameters for experimental and control arm OS data (drawn from 116 trial figures). For α, the value for the control arm was subtracted from the value for the experimental arm. Differences in β were computed by determining the percent change in β in the experimental arm with respect to the control arm (positive values indicate larger β in the experimental arm). Asterisks denote trials that tested immune checkpoint inhibitors. Success in all cases was judged based on the original report and most often corresponded to HR<1 at 95% confidence by Cox proportional hazards regression. Hazard ratio and Weibull ratio of cumulative hazards of four clinical trials in data set: **(B)** NCT01377376 (tivantinib plus erlotinib vs. erlotinib), **(C)** NCT01607957 (TAS-102 vs. placebo), **(D)** NCT00861614 (ipilimumab vs. placebo), **(E)** NCT01673867 (nivolumab vs. docetaxel). The uncertainty in estimating the ratio of cumulative hazards typically falls with time, narrowing the 95% confidence interval depicted in blue.

Fundamental to the model of proportional hazards is the idea that control and experimental arms are described by hazard functions related by a constant of proportionality (the hazard ratio) that does not change over time. From the perspective of Weibull distributions, this means that the two arms have the same shape parameter (i.e. the difference in α values, Δα, is zero); the trial will be successful (hazard ratio < 1) if the experimental arm has significantly larger β value than the control arm. Across 121 comparisons of experimental and control arms from 116 OS trial figures, we found that Δα values actually varied from +0.65 to -0.80 (median absolute value |Δα| = 0.11; **Fig. 6A**). For event-free survival data, 154 comparisons of experimental and control arms from 146 trial figures revealed a range of Δα values from +0.49 to -0.85 (median |Δα| = 0.08) (**Supplementary Fig. S7**). Thus, the assumption in the proportional hazards model that Δα = 0 is frequently violated. To compare this to previous reports of violation of proportional hazards, we used a Grambsch–Therneau test at a significance level of p < 0.1 (Guyot et al., 2012). In our dataset, 18 out of 108 unique publications reporting OS data (∼17%) were found to violate proportional hazards, and a significant deviation from the proportional hazards assumption (Guyot et al., 2012; Rahman et al., 2019) corresponded to |Δα|=0.30. For unique trials reporting event-free survival data, 47/135 (∼35%) were found to violate proportional hazards by the Grambsch–Therneau test at 10% significance, and this significance cutoff also corresponded to |Δα|=0.30. ICI trials tended to violate proportional hazards at a higher rate compared to all other trials (4/5 trials). Thus, differences in Weibull α parameters greater than 0.3 for control and experimental trial arms identify significant deviations from the assumption of proportional hazards (Rahman et al., 2019) and are most common in ICI trials (Alexander et al., 2018; Chen, 2013).

We used parametric fitting to explore the origin and consequence of non-proportional hazards in trial data. Specifically, we used Weibull shape and scale parameters for each trial arm (and confidence intervals thereof) to calculate the ratio of cumulative hazards between experimental and control arms at time *t* (HR_c_(*t*)), for all values of *t* from the start to the end of the trial. This approach returned robust estimates of relative empiric hazard for each trial arm as a function of time without assuming proportionality. Trials that deviated little from the proportional hazards assumption, such as the failed ATTENTION trial (NCT01377376) (Yoshioka et al., 2015) with Δα =0.003 or the successful RECOURSE Trial (NCT01607957) (Mayer et al., 2015) with Δα =0.13, Weibull HR_c_(*t*) (blue lines in **Fig. 6B, C)** closely approximated HR_SP_ from Cox regression (red lines; note that the HR_SP_ under the assumption of proportional hazards is, by definition, time-independent). In the case of the successful trial CheckMate 057 (NCT01673867) (Borghaei et al., 2015) a value Δα =-0.26 denotes a deviation from proportional hazards and we observed that HR_c_(*t*) matched HR_SP_ at only one point in time (**Fig. 6E**). This was also true of CA184-043 (NCT00861614) (Kwon et al., 2014) for which Δα = -0.30; this trial was judged to have failed based on HR_SP_ (**Fig. 6D**). In this trial, the HR_c_(*t*) also matched HR_SP_ from Cox at a single point in time but it fell steadily to <1 by the end of the trial. Unless the shape of the hazard function were to substantially change over one additional month, it seems probable that CA184-043 would have been judged a success by conventional HR_SP_ criteria had it continued slightly longer. Time-dependent success was recently demonstrated in the MK-3475-022/KEYNOTE-022 trial of pembrolizumab with dabrafenib and trametinib for BRAF-mutant melanoma, where the pre-planned analysis at 24 months did not identify a statistically significant benefit (PFS HR of 0.66, 95% CI 0.40-1.07) but a subsequent analysis at a median 36.6 months of follow-up did (PFS HR of 0.53, 95% CI 0.34-0.83) (Ascierto et al., 2019; Ferrucci et al., 2020). We conclude that violations of the assumption of proportional hazards in clinical trial data do not simply involve statistical deviations, but actual variations in treatment effect over time – and this is true of both OS and PFS data. This contrasts the assumption in Cox regression that HR_SP_ has a fixed value over the course of the trial, such that time enters into consideration only insofar as enough events must accrue for HR_SP_ to be judged significantly different from one (Cox, 1972).

To more fully explore the dependence of trial duration on outcome, we simulated a series of trials in which control and experimental arms exhibited a range of differences in α and β values and then plotted the fraction of trials that were successful, as judged by Cox regression under a proportional hazards assumption (HR_SP_ <1 at 95% confidence). In evaluating “success”, we applied Cox regression and ignored violations of proportional hazards as is the currently accepted standard (see Discussion). Success was evaluated at a stopping point defined by 60% of events being recorded (t_A_) or a late stopping point of 95% of events being recorded (t_B_; **Fig. 7; Methods**). For simulated trials in which α was smaller for the experimental than the control arm, the likelihood of success was substantially greater at the late stopping point as compared to the earlier stopping point (**Fig. 7A-B**; conversely, early termination, for apparent futility for example, would incorrectly support a conclusion of inferiority). In these cases, we see that the experimental arm exhibited lower survival at early times and then crossed the control arm at a later time to exhibit higher survival. The greater the value of Δα, the greater the impact of curve crossing and duration of follow-up on outcome. All ICI trials in our data set fall in this category, since their experimental arms have smaller α parameters than their control arms **(Fig. 6A)**, resulting in decreasing HR_c_(*t*) over time and curve crossing. Thus, the time at which these trials are terminated impacts outcome, independent of the number of events needed to reach statistical significance (Mick and Chen, 2015). The reasons for time-dependent therapeutic effects are unknown, but in ICI trials it has been suggested that they are related to initial treatment-related toxicity or delays in treatment effect (Alexander et al., 2018; Chen, 2013; Mick and Chen, 2015; Rahman et al., 2019). We conclude that future ICI trials should consider the impact of time-varying changes in relative hazard (which are detectable via Weibull fitting) on trial success as a factor independent of the number of events needed to reach statistical significance in the estimate of HR_SP_.

**Fig. 7:**
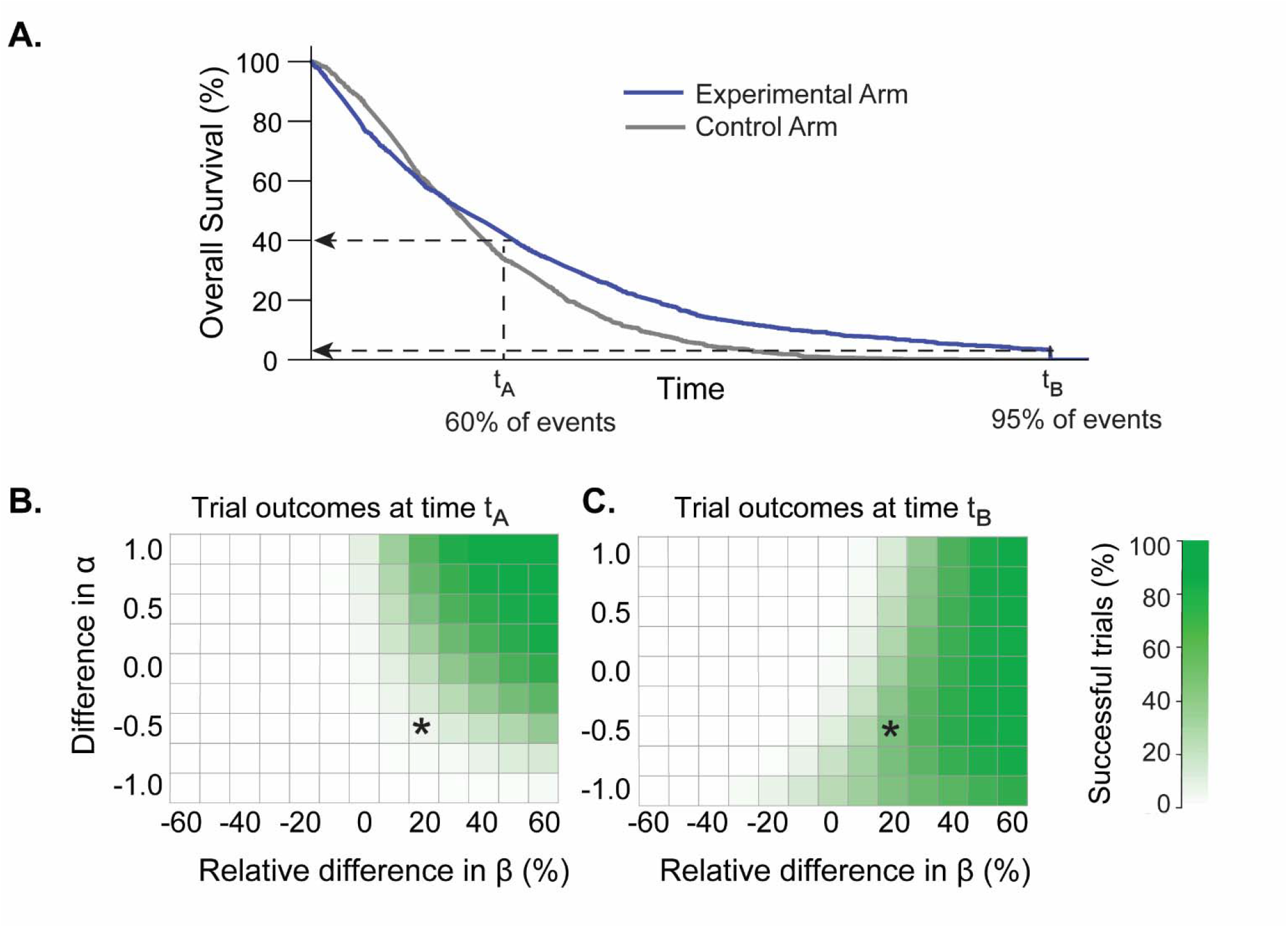
Effect of trial duration on success when proportional hazards is violated. **(A)** One of many simulated trials having a range of Weibull α and β parameters similar to those observed in actual trials reporting OS data; in this trial Δα = –0.5 and the ratio of β for experimental and control arms was 1.2. The labels t_A_ and t_B_ denote times corresponding to 60% or 95% of all trial events (for real OS trial arms in this article, in metastatic cancers, these event rates correspond to median times of t_A_ = 16 months and t_B_= 51 months). **(B)** Percent of simulated successful trials at time t_A_ (left panel) or t_B_ (right panel). The simulated trial depicted in panel A is denoted by an asterisk. “Success” was scored as HR<1 at 95% confidence using the Cox proportional hazards regression; this metric was used despite the violation of proportional hazards because it is the accepted approach for assessing efficacy in pivotal trials (see text for details).

## DISCUSSION

Using a set of ∼220,000 imputed participant survival events from published oncology trials we find that survival functions for solid tumors, including those from trials that report OS or event-free survival data (i.e.: PFS), or are biomarker-stratified, are well fit by two-parameter Weibull distributions. The Weibull α (or shape) parameter defines increasing or decreasing hazard over time and the β parameter is proportional to the median survival time. The excellent fit of survival data to a single parametric function for many types and stages of cancer and across drug classes demonstrates that therapeutic benefit can be well-described by a simple function that models response as varying continuously across a population. In the trials studied here, the likely presence of prognostic factors, or responder and non-responder populations, did not sufficiently separate survival functions to produce bimodal distributions that would have necessitated the systematic use of mixture models or cure-rate parameters.

Our findings support modeling survival in early stage oncology clinical trials by using parametric statistics. Parametric statistics are already used in cost-effectiveness analysis (Hoyle and Henley, 2011) and other simulation studies. Our data uses a large and diverse clinical data set to verify that Weibull distributions are the preferred parametric form and that they have sufficient accuracy for applications in solid tumors. By simulating trials of different sizes, we find that modeling with parametric statistics substantially improves precision with equivalent accuracy: assessing 12-month survival outcomes using a Weibull parametric approach makes a 50-person trial reporting OS data as precise as a 90-person trial evaluated using nonparametric methods. This underestimates the actual benefit of using Weibull forms because nonparametric methods did not return a numerical confidence interval in 20-40% of OS simulated trials with 20-100 patients, whereas parametric estimation was successful in all cases. Thus, the use of parametric statistics should be strongly considered in early phase signal-seeking studies with the goal of rapidly and economically identifying the optimal setting in which to perform phase III trials.

Weibull distributions are also appropriate for cost-effectiveness research for oncology drugs, an increasingly important topic for drug approval in many countries. In this context, it is important to note that Weibull and Log-Normal distributions provide equivalently good fits to IPD and the Weibull form was chosen in the current study because of its interpretability in terms of hazard rates and median survival. Log-Normal distributions may have corresponding advantages in pharmaco-economic analysis (Ouwens et al., 2019). Moreover, insofar as there exist multiple ways to implement parametric statistics, we note that our results pertain specifically to approaches detailed in the **Methods** (these are largely conventional). Alternative statistical approaches to increasing precision, for example by changing significance cut-offs of traditional confidence intervals (to create narrower confidence intervals while still maintaining an acceptable level of Type I error) have not yet been explored in detail.

In current practice, Cox regression is used to compare survival functions based the proportional hazards assumption, which states that the ratio of control and experimental hazard functions is constant over time. Success usually corresponds to HR < 1 at 95% confidence. With respect to Weibull distributions, the assumption of proportional hazards corresponds to no difference in shape, i.e. Δα = 0. However, in real trial OS data, Δα was found to vary from 0.65 to -0.80 with ICI trials as a class having the largest |Δα| values. If we use previously described criteria to determine which violations of proportional hazards are significant (Alexander et al., 2018; Rahman et al., 2019) we find that they correspond to Δα > 0.3 and apply to ∼17% of trials reporting OS and ∼35% of trials reporting PFS data. Analysis of imputed data from published trials, and simulations in which empirical survival functions are resampled, shows that violations of proportional hazards are not statistical curiosities but instead arise from time-varying treatment effects. In the data analyzed here, this was most evident in ICI trials, but was also seen in trials of the BCL-2 inhibitor venetoclax, in which experimental and control arms cross each other some time after trial initiation (Kumar et al., 2020).

The biological basis of time-varying treatment effects (and curve crossing) are not known in detail but could arise from high toxicity in a subset of patients early in treatment, delayed onset of treatment effects, or exceptionally durable responses in some patients (Alexander et al., 2018; Chen, 2013; Mick and Chen, 2015; Rahman et al., 2019). Regardless, the practical consequence is that the duration of a trial has a direct impact on outcome, independent of statistical considerations such as increasing confidence in HR values as trial events accrue (as in Cox regression). We found that it was possible to use Weibull fitting to identify trials judged as failures by Cox regression in which HR was trending steadily below one at the end of a trial but an extension of only a few months was predicted to result in success. We suggest that such considerations be taken into account in the design of future trials, particularly for ICIs.

Additional information that can be mined from IPD to improve future trials includes how sample size and power are estimated. Such calculations are most commonly performed under an assumption of an exponential fit to survival data. Alternative distributions have been proposed for such analyses (Heo et al., 1998; Jiang et al., 2012; Lu et al., 2012; Wu, 2015) but without any means for selecting optimal parameter values for simulation. Using the Weibull fitting described here, empirically-derived parameter values can be drawn directly from past trial data. In some cases, past trial data might also be useful for the generation of synthetic control arms (El Emam et al., 2015). Additional applications include the use of parametric forms for subgroup analysis in phase III and basket trials. Since studies of this type are intended to test therapeutic hypotheses rather than lead to drug registration, the regulatory barriers to using parametric statistics are limited. Parameterized Bayesian trial designs, such as the continual reassessment method (CRM) or escalation with overdose control (EWOC), are other model-based methods already in use used to define specific parameter values and improve the efficiency of phase I studies (Liu and Yuan, 2015).

Even modest improvements in the design and interpretation of oncology clinical trials are likely to have a substantial payoff. The overall approval rate for new oncology drugs remains low: only 3% of drugs tested in a phase I clinical trial and 7% of drugs tested in a phase II study are ultimately found to be superior to standard of care comparators in pivotal phase III studies (Wong et al., 2019). Methods to more accurately understand drug activity in small patient populations are included in the NCI 2020 “provocative questions” (National Institutes of Health, 2019) and could lead to a wider use of master protocol trials (Palmer et al., 2020; Park et al., 2019). Improving trial efficiency and predictability will become increasingly critical as the number of new monotherapies and combinations continues to rise, patient populations become more subdivided based on the molecular characteristics of their tumors, and it becomes impractical to enroll enough patients to test all promising drug treatments (Kolata, 2017).

The use of nonparametric statistics was historically appropriate because treatment effects could be calculated precisely without the need for computation, which was largely infeasible prior to the widespread availability of personal computers (Breslow, 1975; Cox, 1972). Moreover, the proportional hazards assumption appears to be largely valid when scoring OS in the context of cytotoxic chemotherapies (OS trials including chemotherapies in our data set had a median |Δα| = 0.11, below the |Δα| = 0.30 threshold for significant violation). However, the widespread violation of proportional hazards reported in this and previous studies, and its likely origins in the biology of new and more diverse forms of cancer therapy, call for a reconsideration of Cox regression. Several approaches for comparing treatment effects have been proposed including weighted (Fleming and Harrington, 2005) or adaptive log-rank tests (Yang and Prentice, 2010), restricted mean survival test (Royston and Parmar, 2013), and permutation-based approaches (Arfè et al., 2020). It is probably appropriate for regulatory agencies to examine the possible use of such methods in the setting of the ICI trials that are the focus of so much current research.

## Limitations of this study

This work is not a formal meta-analysis or systematic review of a specific treatment regimen or disease, but instead a broadly conceived research study; no treatment decisions should be made based on our findings. A specific limitation of this study is that it uses imputed IPD rather than original data. A related limitation is that we only analyze four tumor types (breast, colorectal, lung, and prostate); extending the analysis to additional cancer types will require imputing IPD from additional trials. Finally, we use Cox regression to determine whether real or simulated trials are “successful” (e.g. **Fig. 6 and 7**) even when underlying survival distributions clearly violate proportional hazards. We do this because a finding of HR <1 at a pre-specified level of confidence is the only widely accepted method for evaluating trial outcomes. Potential limitations in parametric approaches may be addressable by reconstructing a larger number of trials for additional cancer types; however, this is a substantial undertaking.

We are forced to use imputed IPD because original results are simply not released, and most published oncology trial reports do not provide the numerical values used to plot the Kaplan-Meier estimators. Release of numerical data underlying graphical representations has become the norm in pre-clinical research and is at the heart of efforts by funding agencies to make data FAIR (findable, accessible, interoperable and reusable) (Wilkinson et al., 2016). Multiple calls have been made to make IPD from research clinical trials publicly accessible (Danchev et al., 2021; Drazen, 2015) to ensure the reproducibility of study results and facilitate meta-analyses, but compliance remains low. Outside of oncology, calls for reuse of both contemporary and historical control arms have arisen in repurposing trials for COVID-19, particularly when the same set of institutions is conducting many parallel trials outside of a master protocol framework (Karlin-Smith, 2020).

Ongoing data collection efforts relevant to clinical trials include the US National Cancer Institute’s (NCI) Project DataSphere (Hede, 2013), the NCI National Clinical Trials Network (NCTN) and Community Oncology Research Program (NCORP) Data Archive, and The Yale University Open Data Access (YODA) Project (Ross et al., 2018). Unfortunately these projects have substantial limitations with respect to the type of analysis presented here: (i) most IPD are greater than six years old and do not cover many of the drugs of greatest current interest, including ICIs; (ii) most public data derives from control, not experimental treatment arms; (iii) much of the data involves summary statistics not IPD, and requests for underlying data can be strictly limited; (iv) if access to IPD is granted, they are often available online for inspection but are not downloadable for computational analysis. A substantial unmet need therefore exists for primary data from clinical trials to be made available for reuse. One approach is to amend the requirements for data deposition on ClinicalTrials.Gov (per U.S. Public Law 110-85) to include IPD.

## ACKNOWLEDGMENTS

We thank Lorenzo Trippa, Andrea Arfe, Giovanni Parmigiani, Charles Perou, John Higgins, and Michael Kosorok for their helpful comments on this project. We thank Jeremy Muhlich and Zev Ross for their assistance in implementing the cancertrials.io site. We are grateful to all of the patients and investigators who participated in the clinical trials analyzed in this work. This project was supported by NIH grants P50-GM107618 and U54-CA225088 (to P.K.S). D.P. is supported by NIGMS grant T32-GM007753 and F30-CA260780.

## AUTHOR CONTRIBUTIONS

Generating individual participant data set: G.F., B.M.A.

Metadata curation; data analysis: D.P. and A.C.P.

Writing: D.P., A.C.P., and P.K.S.

Manuscript review and editing: D.P., G.F., B.M.A., A.C.P., P.K.S.

Supervision: B.M.A., A.C.P., P.K.S.

Funding: B.M.A. and P.K.S.

## STAR METHODS

### Resource availability

#### Lead Contact

- Further information and requests should be directed to and will be fulfilled by the lead contact, Peter K. Sorger (; cc).

#### Materials Availability

- This study did not generate new unique reagents.

#### Data and code availability

- The article includes all datasets and code generated or analyzed during this study.

### Method details

#### Individual participant data imputation and curation

The original data set consisted of 152 unique trials in breast, colorectal, lung, and prostate cancer in the metastatic and non-metastatic settings from 2014-2016. Trials were removed from the original data set if there were any inconsistencies in the imputed patient data as compared to its associated clinical trial (e.g.: differing numbers of patients from the publication at-risk table and imputed data). The quality of the data imputation was confirmed quantitatively, by calculating the hazard ratio for imputed data and comparing it to the corresponding trial’s reported hazard ratio, and qualitatively, by overlaying the Kaplan-Meier curve generated from the imputed data on top of the published curve. Trials with a hazard ratio difference greater than 0.1, or with perceptible visual differences, were removed from the final data set and not analyzed further (**Supplementary Data File S1**).

#### Parametric fitting of patient survival data

The event times for the imputed patients, either death for overall survival distributions or surrogate events in the case of event-free survival distributions, were compared to the event times simulated under each parametric distribution. The likelihood of a specific parametric form to fit patient data was computed by maximum likelihood estimation. Specifically, the relative likelihood of a patient event taking place at a particular point in time was calculated under that parametric distribution’s probability density function. The likelihood of a censoring event taking place was calculated by integrating the probability density function (the cumulative density function), and computing the likelihood of a patient event taking place in the trial after the censoring time (1-the probability at that time under the cumulative density function). This procedure was repeated for all patient events in an arm of a clinical trial, and the overall likelihood of a fit was calculated by multiplying all relative likelihoods.

#### Computing R^2^ explained by the Weibull fit

For imputed patient events in a clinical trial arm, the event times (deaths or surrogate events) and corresponding percent survival (OS or event-free survival) were computed. Weibull parametric fitting was used to obtain the best-fit α and β values corresponding to the imputed patient data. The differences between the survival distribution under a best-fit Weibull model and the imputed data were analyzed through a Weibull plot (Nelson, 2004). In this approach, the event times and corresponding survival are normalized such that if the data follow a Weibull distribution, the points will be linear. The event times were normalized through the transformation: ln t/β, while survival was normalized by: ln (-ln S (t))/ α. Coefficient of determination (R^2^) values were calculated to determine the goodness of Weibull fitting for all trial arms in the data set.

#### Computing Weibull fits to trials of immune checkpoint inhibitors

Trials of immune checkpoint inhibitors were selected from the data set (five in total). Each trial’s OS and PFS IPD were fit to a 1) single Weibull distribution 2) a mixture distribution made of two Weibull distributions. The quality of all fits was assessed by computing the variance in event time explained by the fit and the Akaike information criterion (AIC)(Akaike et al., 1973). An additional set of simulations was performed to account for the periodicity of radiological scans in detecting progression events, and the quality of fit was quantified by the coefficient of determination (R^2^).

#### Assessing relationship between trial scan time and PFS drop

Trials of immune checkpoint inhibitors in oncology were obtained through a PubMed search of the terms “neoplasms” or “cancer” and “Clinical Trial, Phase III” along with therapies of interest (“ipilimumab” or “pembrolizumab” or “nivolumab”). The search was filtered to yield 25 trials with PFS data and a reported scan time, in addition to the five trials in the original data set, for a total of 30 trials for subsequent analysis. PFS curves were extracted from each of the trials and images were analyzed using DigitizeIt software (Rakap et al., 2016) (Braunschweig, Germany) to estimate the timing of the PFS drop in each survival curve. The trial scan time interval was obtained from each publication’s methods, blinded from the image associated with each trial. All extracted values can be found in **Supplementary Data File S3**.

#### Simulating differences in trial success based on α and β values

Control and experimental arms of clinical trials were simulated 1000 times by drawing 100 patient events from Weibull distributions with differing α and β values. α values in the experimental arm ranged from 0.5 to 4.5, β values from 0.4 to 1.6, and control arm parameter values were kept constant (1.5 and 1 respectively); Figure 7 shows results in the region of interest, from experimental arm α=0.5 to 2.5. Events were censored at either early time points (corresponding to a ∼60% event rate, a time equal to the control arm β value) or later time points (corresponding to a ∼95% event rate, a time equal to four times the control arm β value). Trial success was calculated for each simulation by using a Cox regression at a significance level of p=0.05, in accordance with standard statistical methods used in clinical trials.

#### Calculating the precision and accuracy of parametric and nonparametric survival estimates across sample sizes through subsampling

IPD for each trial arm in the data set was extracted. 213 OS and 273 event-free survival trial arms were used for further analysis; these trial arms had at least 100 patients (the maximum number of patient subsampling events used in this experiment) and at least one event (i.e.: death, progression) taking place before 12 months. For each trial arm, 20-100 patient events (with a step size of 10 events) were subsampled from the imputed IPD. At least 3 non-censoring events were selected during each sampling simulation. This procedure was repeated ten times per sample size and trial. Parametric and nonparametric 95% confidence intervals for 12-month survival were computed for every sampling simulation.

Accuracy and precision plots were constructed for the subset of simulated trial arms returning numerical nonparametric confidence intervals (125 OS trial arms and 99 event-free survival trial arms). Note that nonparametric estimates did not return a numerical confidence interval for 41% of OS trial arms and 64% of event-free survival trial arms, while Weibull fitting made it possible to calculate 12-month confidence intervals for every trial arm in every simulation.

#### Quantification and statistical analysis

All analysis was performed using Wolfram Mathematica Version 12.1.0.0. Details of the statistical analysis performed, exact values of n and what they represent, definitions of the summary statistics used, definitions of significance, and trial inclusion and exclusion criteria can be found in the Method Details, Figure captions, and Results sections of the manuscript.

## SUPPLEMENTAL INFORMATION TITLES AND LEGENDS

**Supplementary Fig. S1.**
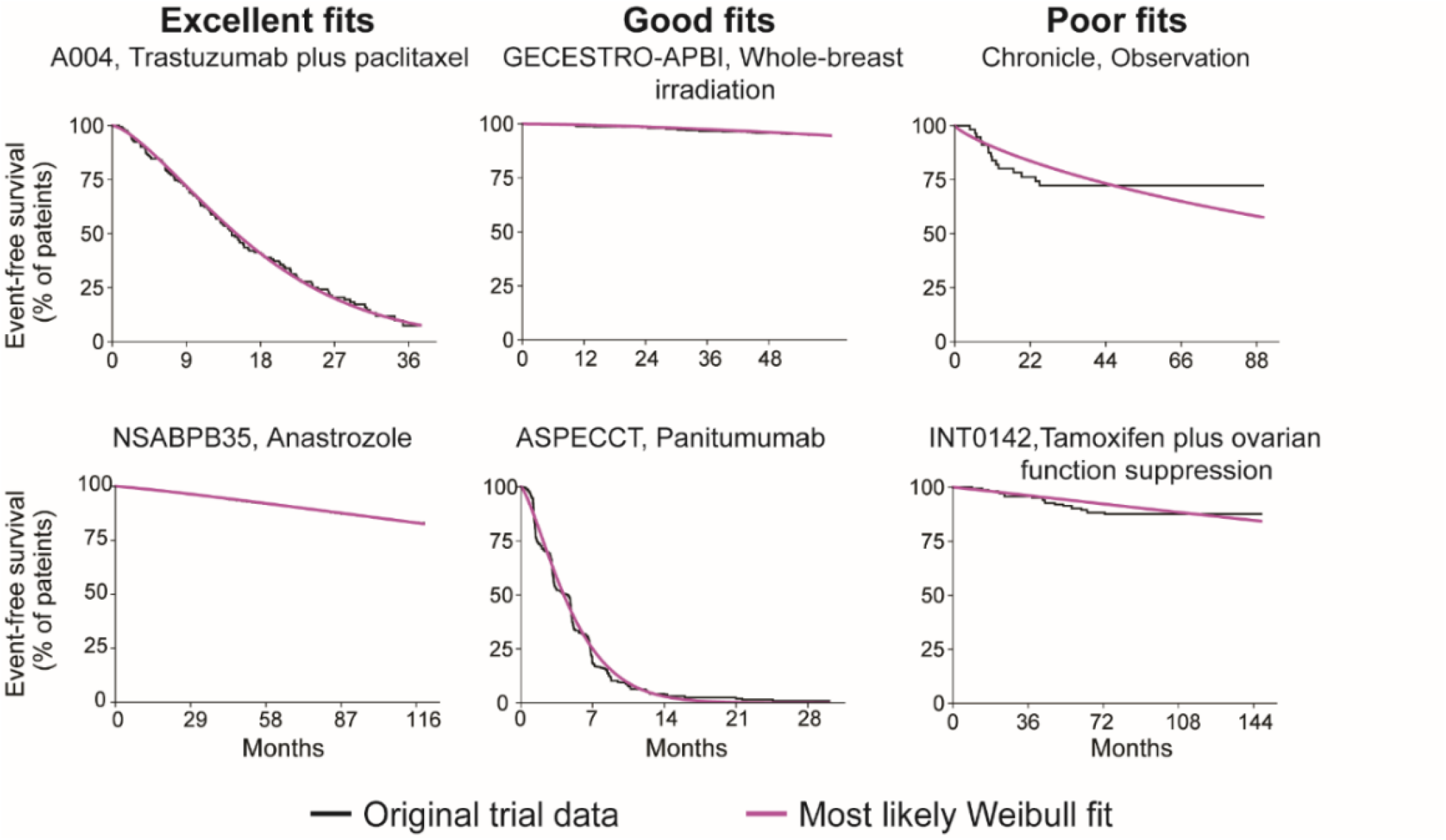
Related to Figure 2: Representative fits of Weibull distributions to event-free survival data. Weibull fits to data for plots of the Kaplan-Meier estimator falling in the top 25^th^ percentile quality for all fits (excellent fits; NCT00294996, NCT00053898), at the 50^th^ percentile (good fits; NCT00402519, NCT01001377), and in the bottom 25^th^ percentile (poor fits; NCT00427713, NCT number not reported). Data derived from trials reporting event-free survival data, primarily PFS (301 survival curves from 146 figures).

**Supplementary Fig. S2.**
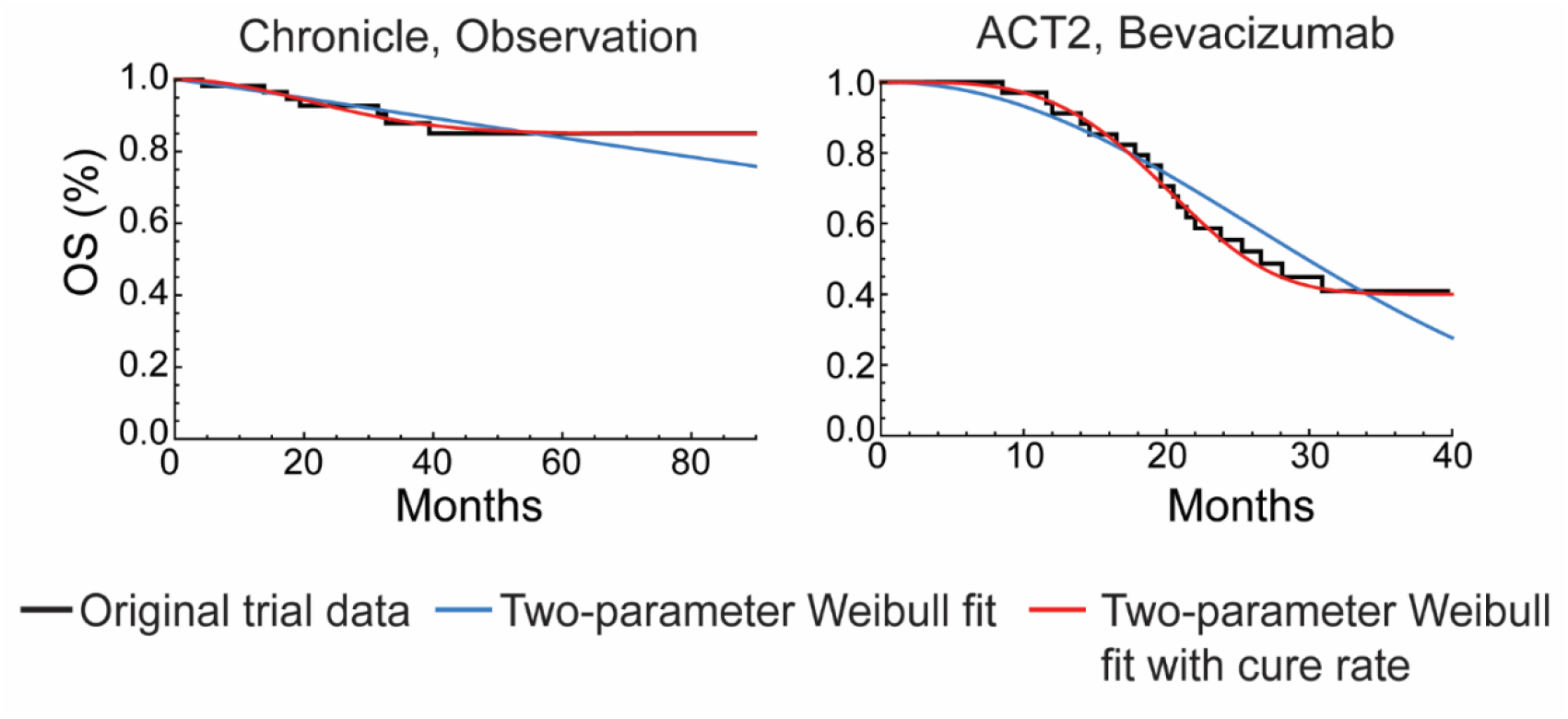
Related to Figure 2: Improvements in fit of Weibull distributions to overall survival data using three-parameter models. Two-parameter fits for the trial arms in Figure 2 representative of the bottom quartile of all Weibull fits (NCT00427713, NCT01229813) and two-parameter fits with an additional cure rate parameter.

**Supplementary Fig. S3.**
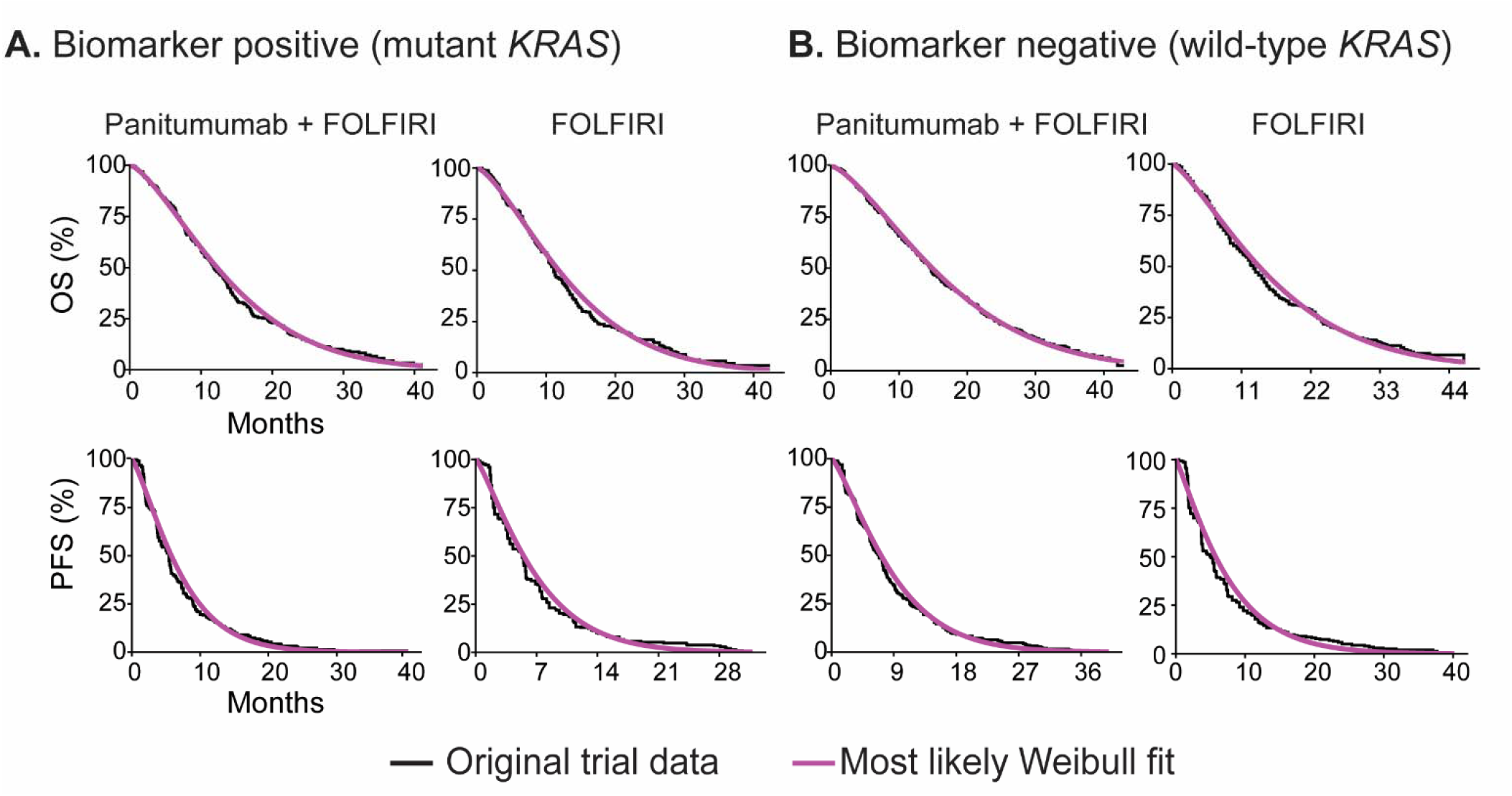
Related to Figure 2: Fit of Weibull models to overall survival and progression-free survival data from biomarker stratified trials. Data obtained from the 20050181 trial (NCT number not reported).

**Supplementary Fig. S4.**
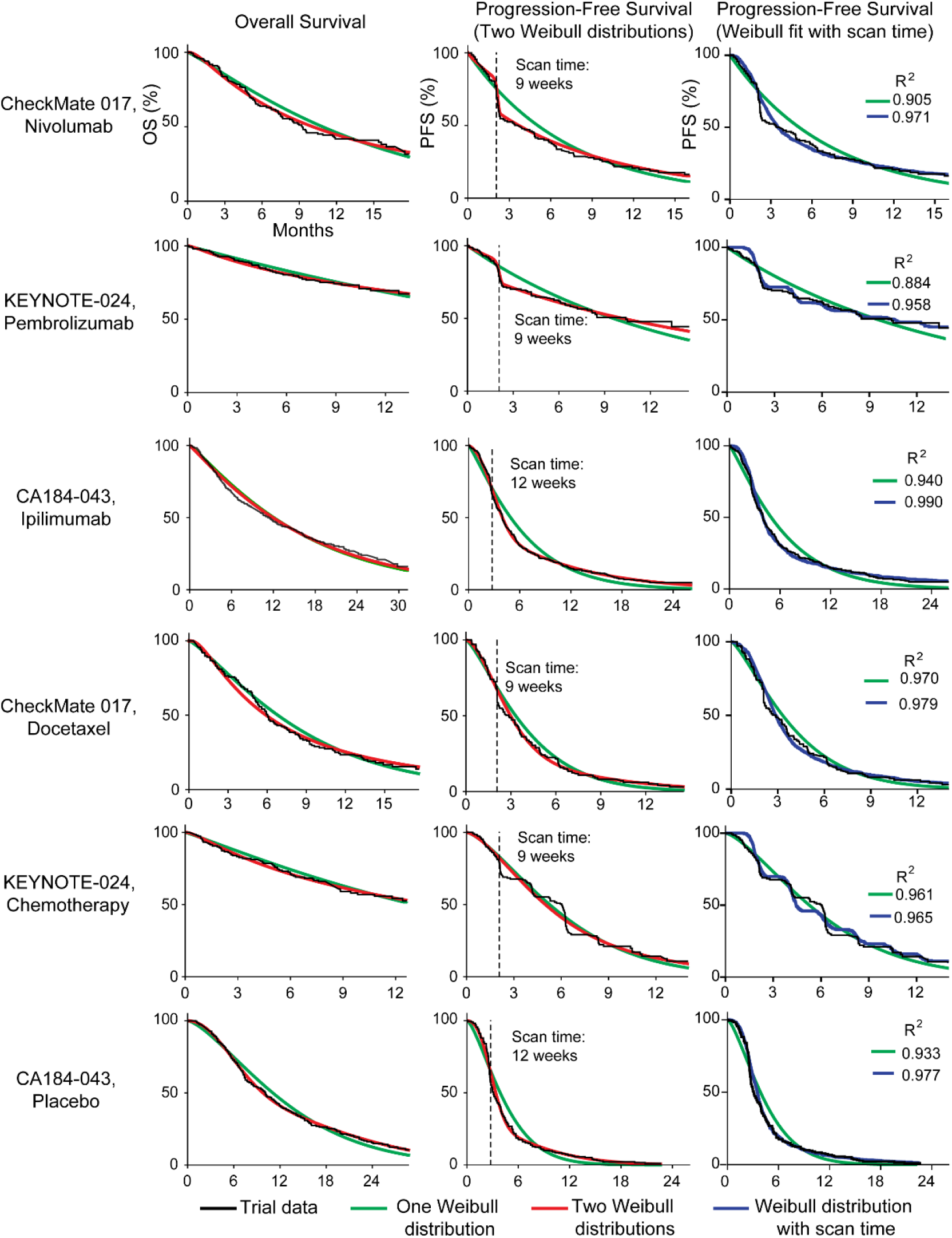
Related to Figure 3: Additional fits to overall survival and progression-free survival data for trials of immune checkpoint inhibitors. Progression-free survival and overall survival distributions for three trials of immune checkpoint inhibitors with one and two-distribution fits using Weibull functions, as well as simulations that account for the periodicity of radiological scans to detect progression (NCT01642004, NCT02142738, NCT00861614).

**Supplementary Fig. S5.**
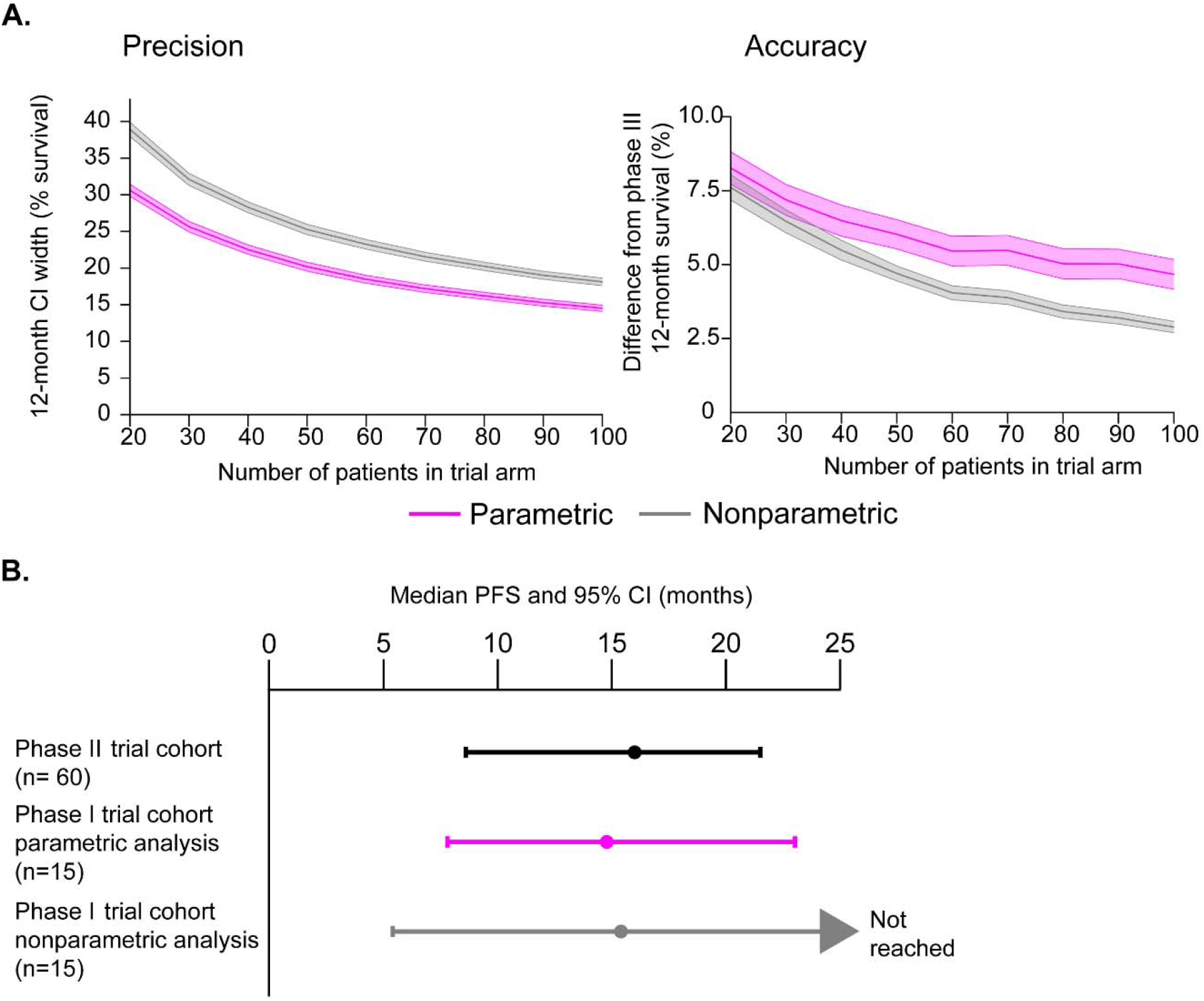
Related to Figure 4: Impact of parametric statistics on precision and accuracy of event-free survival estimates. **(A)** Comparison of nonparametric and parametric (Weibull distribution) methods to compute 12-month event-free survival confidence intervals for trials with small cohorts (20 to 100 patients) produced by randomly subsampling patient events from 99 event-free survival trial arms. Note that nonparametric estimates did not return a numerical confidence interval for 64% of event-free survival curves, while Weibull fitting made it possible to calculate 12-month confidence intervals for every survival curve in every simulation. Precision is defined as the width of the confidence interval in percent survival. Accuracy is defined as the absolute difference between the 12-month survival estimated from small cohorts (for each of 10 simulations per sample size and trial), and the value computed from all patients in a given phase III trial. Shaded regions are 95% confidence intervals. **(B)** Median PFS confidence intervals calculated with parametric and nonparametric methods on phase I clinical trial data, and the corresponding phase II study results (MK-3475-022/KEYNOTE-022; NCT02130466).

**Supplementary Fig. S6.**
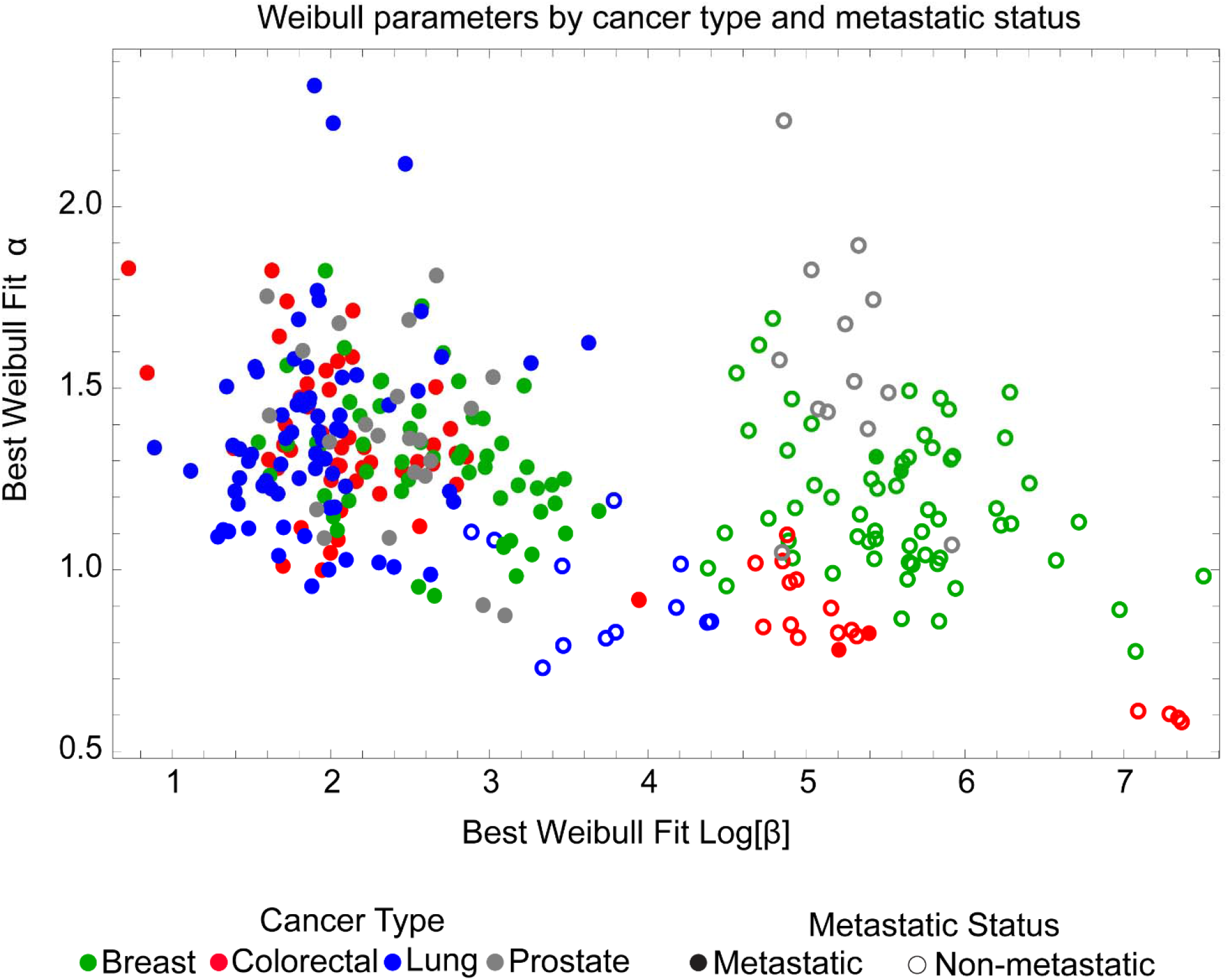
Related to Figure 5: Best fit Weibull parameter values for trials reporting event-free survival data. Weibull fits for event-free survival curves labeled by metastatic status and cancer type (encompassing 301 survival curves from 146 trial figures).

**Supplementary Fig. S7.**
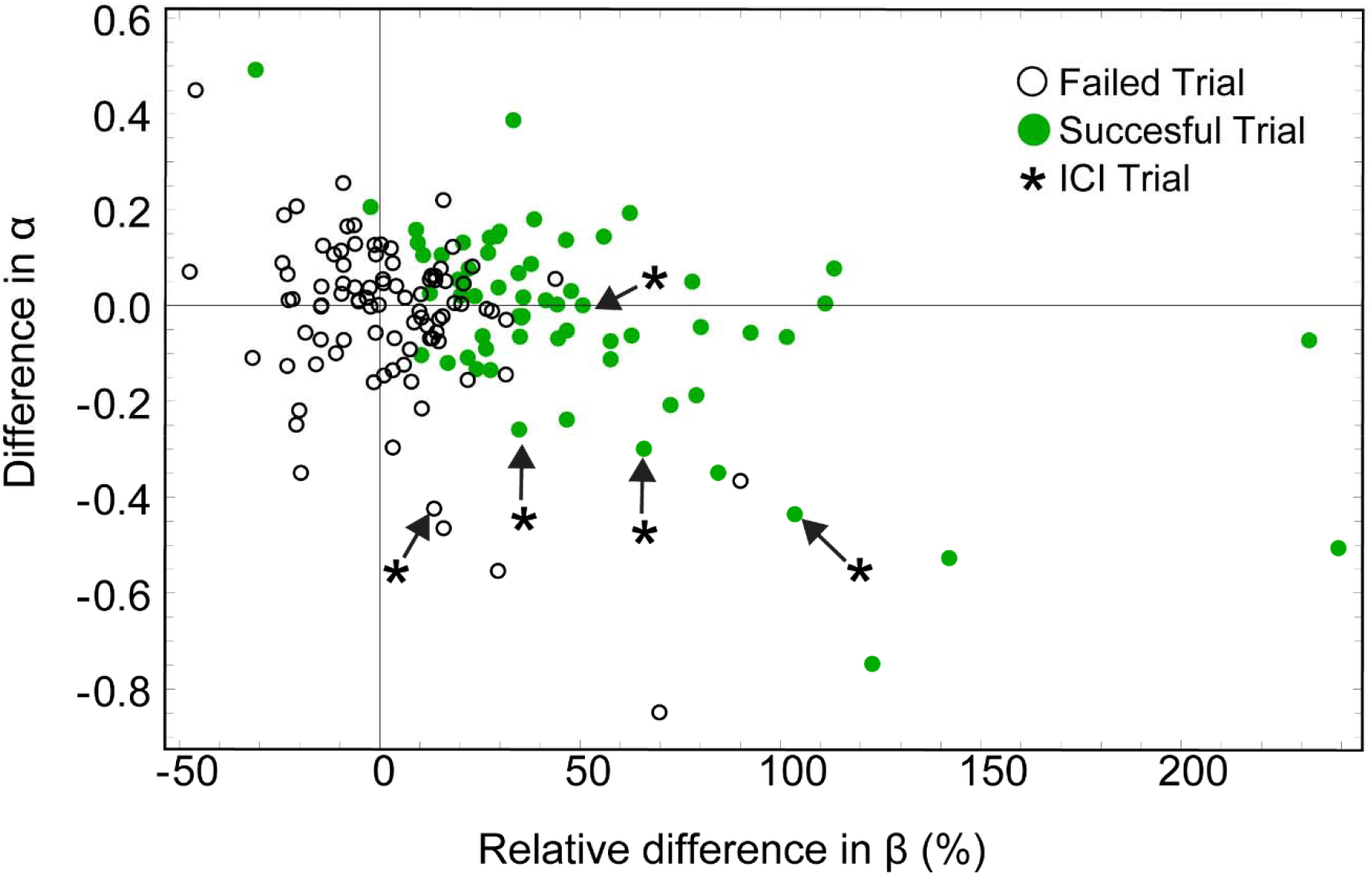
Related to Figure 6: Parameter values for Weibull fits to event-free survival data scored by outcome of the trial. For α, the value in the control arm was subtracted from the value for the experimental arm. Differences in β were computed by determining the percent change in β value in the experimental arm with respect to the control arm (positive values indicate larger β in the experimental arm). Success in all cases was judged based on the original report and most often corresponded to HR<1 at 95% confidence by Cox proportional hazards regression.

**Supplementary Data File S1.** Clinical trial metadata and IPD. Trial metadata file includes: trial name, author, registration number, journal, publication date, cancer type, cancer metastatic status, whether a significant difference was found between the trial experimental and control arm, treatment name, treatment type, and number of patients enrolled in the trial. Comparisons between the imputed trials’ hazard ratios and the original trial hazard ratios are included to assess imputation quality (procedure described in Methods). IPD is provided as 262 .csv files. Each .csv file contains IPD from a different figure from a published clinical trial. Description of all variables included in metadata and .csv files can be found in “README.txt”.

**Supplementary Data File S2.** Analysis code. Each piece of code is provided in a folder containing a Mathematica Notebook (.nb), all data required by the code, and the corresponding code output. With source data kept within the same folder as the code, the Mathematica Notebook can be executed in Wolfram Mathematica version 11 by selecting “Evaluate Notebook” from the “Evaluation” menu.

**Supplementary Data File S3**. Weibull fitting of ICI trial arms. The first tab contains percent variance explained and AIC for one and two-distribution fits of ICI trial arms. The second tab contains the timing of ICI trial PFS drops and the corresponding trial scan times.

